# Neuronal communication subspaces arise across prefrontal-striatal circuits prior to gaze-directed object selection and develop with learning

**DOI:** 10.64898/2026.11.24.768911

**Authors:** Peter G. Poggi, Seyed-Alireza Hassani, Adam Neumann, David Brust, Kianoush Banaie Boroujeni, Paul Tiesinga, Thilo Womelsdorf

**Affiliations:** Department of Psychology, Vanderbilt University; Nashville, TN, USA; National Institute of Neurological Disorders and Stroke, National Institutes of Health; Bethesda, MD, USA; Donders Institute, Radboud University; Nijmegen 6525 EN, The Netherlands; Vanderbilt Brain Institute; Nashville, TN, USA; Department of Biomedical Engineering, Vanderbilt University; Nashville, TN, USA

## Abstract

How neuronal ensembles determine the most relevant object to attend to remains unresolved but likely involves coordinating information across multiple brain areas. We found support for this hypothesis in latent firing co-variations across the prefrontal cortex (PFC), anterior cingulate cortex (ACC), and head of the caudate (CD) during and after attention-set learning. Before gaze shifted to an object, neuronal communication subspaces emerged across area pairs. While exploring options during learning, ACC and PFC led communication over the CD prior to gaze shifts, which switched to the CD and PFC leading the ACC when exploiting learned attention sets. Neurons participating in exploration- and exploitation-specific subspaces were partly segregated and carried information about the value and location of the to-be-selected objects, suggesting they guide attention and gaze.

## Main Text

Flexible behavior requires learning what is relevant and using that knowledge to guide attention and action. As behavior shifts from exploring options with uncertain values to exploiting learned associations, distributed brain areas are involved in selecting visual objects, making choices, evaluating outcomes and updating behavioral policy (*1*). How this occurs in neural networks remains poorly understood. However, advances in large-scale recordings and population analysis methods have made it possible to study multi-area coordination at the level of neuronal populations (*2*–*7*). These approaches have revealed that interactions between areas can be concentrated within low-dimensional patterns of population activity, termed communication subspaces, in motor, visual, and higher-order networks (*8*–*13*). Whether and how such communication subspaces arise and change during and after learning of attention sets remains unknown.

The dorsolateral prefrontal cortex (PFC), the anterior cingulate cortex (ACC), and the head of the caudate (CD) are each key components of the neural network responsible for flexible learning. During learning, the PFC has been shown to encode goal-directed representations and task context (*14*). The ACC motivates information seeking to resolve uncertainty and monitors for violations of expectations (*15*–*17*), and the CD encodes value, supports motor actions, and enhances learning (*18*–*20*). While neuronal populations in these areas can form local low-dimensional firing patterns (*21*–*24*) and show brief periods of correlated firing (*25*), it has remained unknown whether PFC, ACC, and CD engage in communication subspaces to coordinate information. Here, we addressed this question and found that active neuronal communication subspaces emerged during visual object selection across the PFC-ACC-CD network while nonhuman primates first explored the value of visual objects and then exploited learned attention sets. Neurons were partially segregated into exploration- or exploitation-specific subnetworks and conveyed the value and location information needed to guide attention and choice (*26*).

### Learning attention sets involves transitioning from exploration to exploitation states

We recorded activity from 5,088 well-isolated single units (1,167 in the PFC, 890 in the ACC, and 3,031 in the CD) in two adult rhesus macaque monkeys performing a feature learning and set-shifting task (**Fig. 1A**; **Fig. S1**). The task involved using gaze to select one of three objects that differed in features of two feature dimensions (e.g. body shapes and arm styles). On a given trial the subjects freely sampled objects until selecting one by fixating it for 0.7 s. If fixation was broken between 0.3 s and 0.7 s, the trial was aborted. After selecting an object, the subjects were probabilistically rewarded (p = 0.85) by gaining tokens for correct choices or losing tokens for incorrect choices. Tokens filled a token bar that cashed out for fluid reward when filled (**Fig. 1B**). In each recording session, target features changed across 36 blocks of ∼30-60 trials each. Block changes were uncued, requiring the subjects to update their beliefs and learn by trial and error whatthe newly rewarded target feature was. The rewarded target feature within a block was either a distractor feature from the previous block, requiring set shifting, or a novel feature, requiring learning a new feature-reward set. To control for motivational variables, blocks also differed in the incentives for correct choices (+3 or +2 token gains) and the incurred loss for incorrect choices (-3 or -1 tokens) (**Fig. 1B**).

**Fig. 1:**
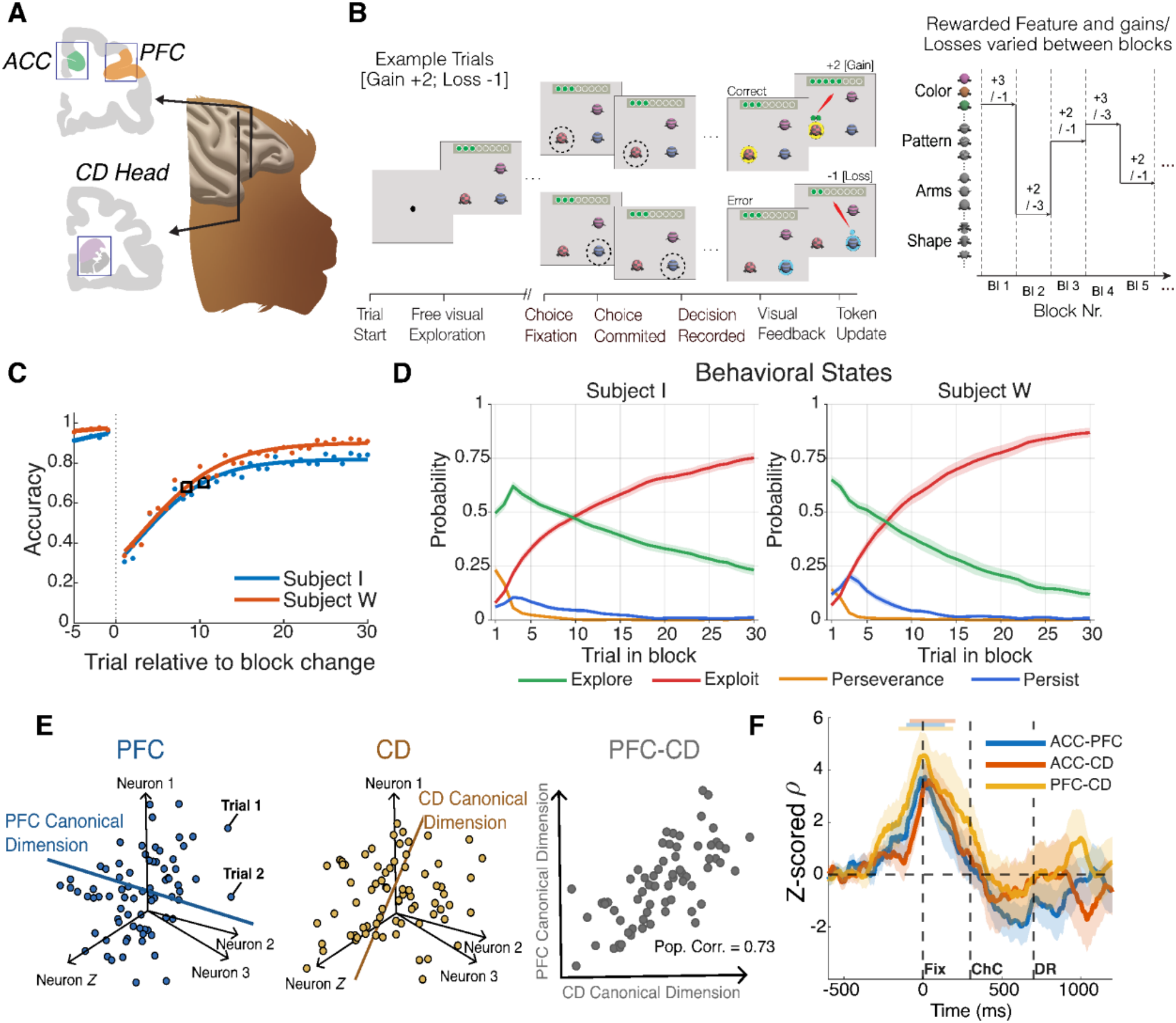
Task paradigm and methods. (**A**) Anatomical diagram of recording sites in the ACC, Brodmann area 9/46 of the dlPFC, and the head of the CD. (**B**) Example trial schema. Trials were initiated with a blank screen followed by the onset of quaddles. Subjects were free to visually explore for up to 5 s before making a choice by fixating an object for .7 s; if they broke fixation after .3 s, the trial was aborted. After the decision was recorded, subjects received visual feedback in the form of a halo, and the token bar was incremented. There were 36 blocks per session with reward conditions varying randomly between blocks. (**C)** Overall learning curves of both subjects with the average trial to criterion (80% accuracy in 10 prospective trials) delineated by black boxes. (**D**) Probabilities of each subject occupying a certain behavioral state determined from the hidden Markov model (HMM) for each trial. (**E**) Schematic of CCA canonical dimensions in neural space maximizing inter-areal correlation. After Semedo et al. (**F**) Zero-lag CCA revealed peaks in inter-areal alignment around fixation (colored bars are p < .05 cluster sign-flip test).

Subjects learned target features quickly, reaching criterion (80% accuracy in prospective 10-trial window) at 9.2 ± 0.9 (Subject W) and 11.6 ± 0.8 (Subject I) (mean ± s.e.m.) trials within a block, and reaching a mean accuracy of 97% (Subject W) and 93% (Subject I) in the last 5 trials of the block (**Fig. 1C**). Criterion performance was reached faster when subjects learned new targets than when reassigning a previous distractor feature to be a new target, while criterion performance was reached similarly in different motivational contexts (**Fig. S2**). We determined for each trial whether the choice reflected exploring an object to learn about its value, or exploiting an object’s learned value by choosing it consistently using a hidden Markov model (HMM) (*27*). The HMM accounted for behavior using four separable latent states, which we labeled exploration (choosing a new object), exploitation (choosing the target object repeatedly), perseverance (choosing a rewarded target from a previous block), and persistence (repeatedly choosing a distractor) (*see* **Suppl. Methods**). During the first few trials of the block, subjects primarily occupied the exploration state, with some trials also belonging to perseverative and persistent states. Consistent with the number of trials needed on average to reach criterion performance, the probability of exploration versus exploitation states transitioned at trial 7.2 ± 0.7 (Subject W) and 9.2 ± 0.9 (Subject I) (**Fig. 1D**).

### Inter-areal connectivity increases within neuronal communication subspaces during object selection

To understand how the PFC, ACC, and CD coordinate during the task, we used canonical correlation analysis (CCA) to analyze active neuronal communication subspaces (*28*). Here we define an active neuronal communication subspace as a low-dimensional latent space with above-baseline population-level co-fluctuations of neural activity between areas after accounting for average firing rates and within-area correlations. Given a high-dimensional neural space with the activity of each neuron as an axis, the activity at a given time point *t* on a given trial is a point in this neural space. CCA finds directions (canonical components) across trials in neural space that maximize the correlation between areas after removing within-area correlations (**Fig. 1E**; *see* **Suppl. Methods**). We used the correlation of the first canonical components (*ρ*) as our measure of the magnitude of inter-areal connectivity within communication subspaces (hereafter subspace magnitude) following previous work (*11, 13*). We found apparent CCA peaks around the time when subjects shifted gaze to the chosen object across multiple sessions (**Fig. 1F**). This peak was visible and above baseline across all area pairs (sign-flip cluster permutation test p < .05).

### Fronto-striatal subspaces increase from explore to exploit and precede object selection

After finding communication subspaces around fixation-based object selection, we hypothesized that learning attention sets would modulate the strength, latency, and directionality of inter-areal connectivity within them, resulting in stronger connectivity when transitioning from exploring objects during learning to exploiting learned top-down feature-reward associations. To test this, we performed time-lagged CCA separately on exploratory and exploitative trials around the time of fixation. Time-lagged CCA infers the directionality of inter-areal connectivity; if *ρ* is higher when neural activity in area *X* is compared with future neural activity in area *Y* versus concurrent activity, the net direction of information flow goes from area *X* to *Y* and vice versa (*11*).

First, we found that communication subspaces emerged around fixation onset during both learning and exploration and increased in magnitude after learning across the ACC-CD and ACC-PFC pairs (cluster permutation tests p < .05) (**Fig. 2A**). For PFC-CD neural ensembles, subspace magnitude was similarly high in both explore and exploit trials, and it remained high for longer during exploration than exploitation (cluster permutation test p < .05) (**Fig. 2A**). Second, inter-areal communication subspaces arose before fixation onset for all area pairs in both explore and exploit, except for ACC-CD ensembles during the exploration state (Holm-adjusted sign-flip p < .05) (**Fig S3B**). Mean latencies were -62 ms, -102 ms, -142 ms relative to fixation onset during exploration, and -72 ms, -202 ms, -307 ms during exploitation for ACC-CD, ACC-PFC, and PFC-CD subspaces, respectively (**Fig. S3A**). These results show communication subspaces preceded fixation onsets and were not a consequence of them.

**Fig. 2:**
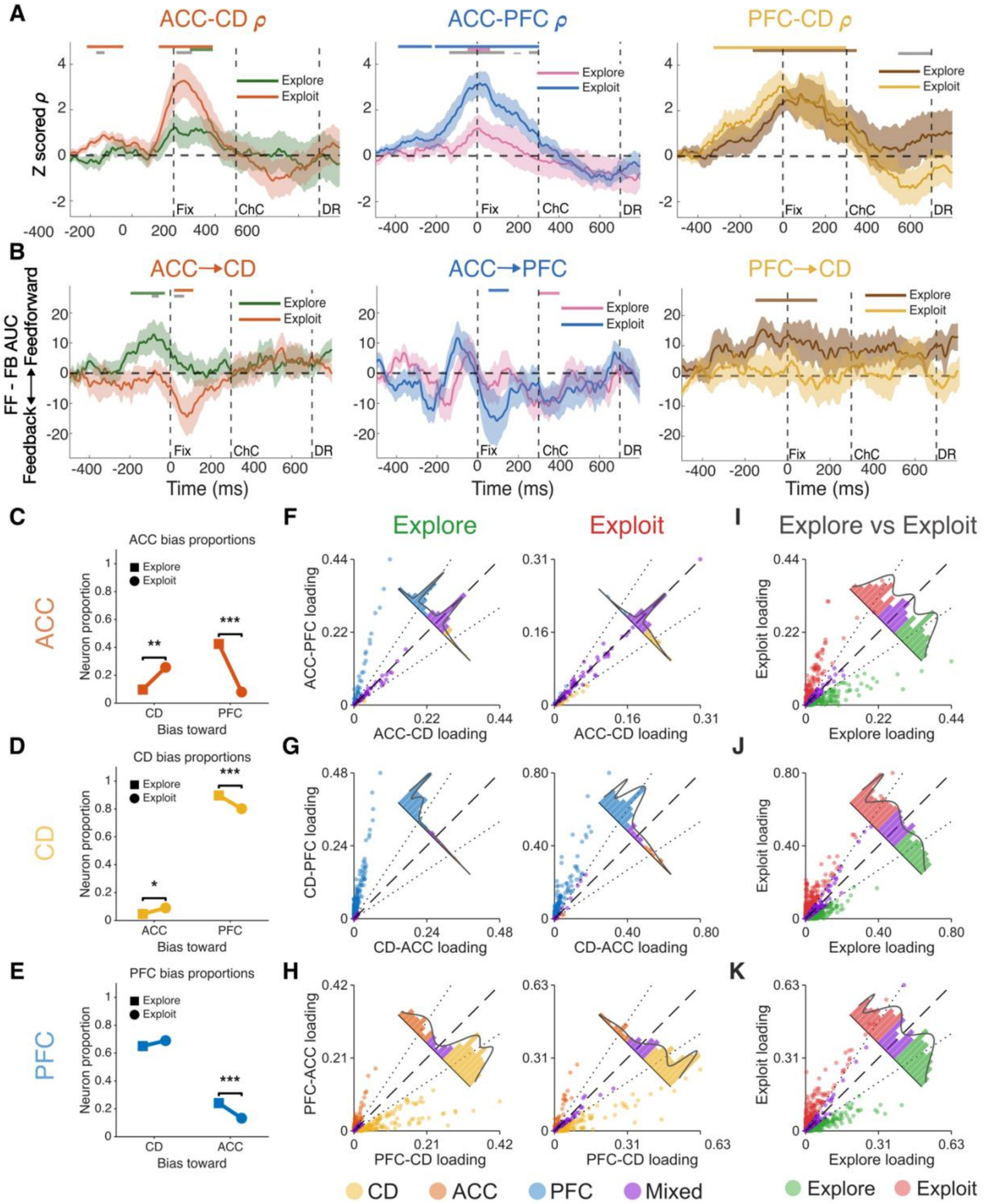
Communication subspace magnitude, direction, and neuronal composition change in explore and exploit. (**A**) Average time-lagged CCA *ρ* during explore and exploit trials across pairs (colored bars p < .05 baseline cluster sign-flip test, grey bars p < .05 between area cluster permutation test). (**B**) Directionality of inter-areal connectivity derived from subtracting positive delay correlations from negative ones (colored and grey bars same as above). (**C**) Proportion of ACC neurons biased towards each other area during explore and exploit. (**D** and **E**) Same as above but for CD and PFC. (**F, G, H**) CCA loadings of each neuron in an area plotted across area pairs along with Gaussian mixture models. (**I, J, K**) Loadings of each neuron plotted for explore and exploit.

We next tested how the strength and latency of subspaces varied across trials relative to the transition from explore to exploit states. Subspace strength gradually increased from explore to exploit for ACC-CD and for PFC-CD ensembles (for each: permutation test p < .05) and tended to increase for ACC-PFC ensembles (permutation test p < .1) (**Fig. S4A**). The peak strength of the PFC-CD subspaces emerged in first trial after the explore to exploit transition and three trials after the transition for ACC-CD and ACC-PFC ensembles. Subspace peaks were higher after learning for the ACC-PFC and ACC-CD pairs in the New condition (**Fig. S5**) and were stronger for ACC-PFC ensembles in exploit states when motivational incentives were highest and punishments were lowest (**Fig. S6**). We also observed larger CCA peaks in correct versus error trials across all area pairs (**Fig. S7)**.

We then used time-lagged CCA to assess lead-lag relationships between areas. During exploration, the ACC and the PFC led over CD (cluster permutation sign-flip test p < .05) (**Fig. 2B**). This ACC / PFC lead over the CD emerged when the inter-areal connectivity started to rise and was evident before gaze shifted to the object (cluster permutation sign-flip test p < .05) (**Fig. 2B**). These pre-fixational directional effects vanished after exploration transitioned to exploitation. After learning, activity in the PFC and the CD led the ACC, but this net directional effected was evident after fixation onset (**Fig. 2B**).

### Separate groups of neurons participate in exploration or exploitation subspaces

Neuronal communication subspaces reflect the co-fluctuations of ensembles of individual neurons. Neurons may participate in communication subspaces with only one other area, forming bi-area interactions, or with two areas, forming a tri-area subspace. We assessed this by extracting the subspace loadings of the individual neurons around the time of fixation onset with each of the other areas and quantifying whether loadings were similar across areas or skewed towards one of the areas (*29, 30*) (**Fig. 2C,D,E**). We found that individual neurons fell into different groups, participating in either two-area or three-areas subspaces, evident in trimodal distributions that were best fit with 3 versus 2 or 1 Gaussians (Lilliefors test for normality, p < .001) (**Fig. 2F,G,H**).

ACC neurons were more likely to participate in a PFC biased subspace during exploration than exploitation (42% during explore versus 8% during exploit). This finding was paralleled by more PFC neurons participating in PFC-ACC subspaces during explore than exploit states (24% / 13% during exploration / exploitation, Holm-adjusted McNemar test p < .001). The opposite pattern was evident for the ACC-CD pair with more ACC neurons participating in subspaces with the CD during exploitation than exploration (10% during explore versus 26% during exploit) (Holm-adjusted McNemar test p < .01) (**Fig. 2C,F**).

While ACC-PFC ensemble participation was larger during exploration, this was not true for PFC-CD ensembles. Similar proportions of PFC neurons participated in PFC-CD subspaces during explore and exploit states (65% / 69 % during exploration / exploitation, no significant difference) (**Fig. 2D,G**), and while a large proportion of neurons in CD participated in CD-PFC subspace ensembles during exploration (89%), fewer participated during exploitation (80%) (Holm-adjusted McNemar test p < .001) (**Fig. 2E,H**). A small population of CD neurons participated in subspaces with the ACC overall, but the proportion increased after learning (5% / 9 % during exploration / exploitation, Holm adjusted McNemar test p < .05) (**Fig. 2E,H**).

Together, these results showed that the proportion of neurons participating in subspaces during explore and exploit differed prominently across exploration and exploitation within inter-areal subspaces. To discern how neurons participated differentially during and after learning, we compared their loadings in explore versus exploit communication subspaces. The majority of neurons in all three areas showed loadings for subspaces preferentially either during the explore or the exploit states but not both states with 76% / 81% / 78% of neurons in ACC / PFC / CD participating only in explore or in exploit subspaces but not in both states (Holm-adjusted one-sided sign test p < .001 for all areas) (**Fig. 2I,J,K**).

### Putative PFC interneurons contribute to the PFC-CD subspace during learning

Communication subspaces could be supported differentially by pyramidal neurons and interneurons. Given the prominent role of interneurons in gating information flow (*31, 32*) and their heightened engagement during learning in the PFC and CD (*33, 34*), we hypothesized that they might participate more strongly in communication subspaces. Our datasets distinguished nine distinct neuron clusters based on their action potential dynamics (**Fig. 3A,B and Fig. S8**) (*35*). Two of those clusters were composed of narrow-spiking neurons indicative of prefrontal inhibitory interneurons (*36*), while seven clusters contained broad spiking neurons (**Fig. 3C**). To test which neurons contributed differentially to inter-areal connectivity, we performed an omnibus test on all neuron clusters and quantified the observed versus expected rates of high loading neurons (median split) across all loading distributions (*see* **Suppl. Methods**). We found that only one—cluster 8 neurons from the PFC (**Fig. 3D**)—was disproportionately represented (permutation omnibus p = 0.00343) (**Fig. 3E**), and this was one of the two narrow-spiking clusters. This putative interneuron cluster included more high loading neurons than expected during exploration states, and specifically in the PFC-CD communication subspace (permutation omnibus p = 0.00343) (**Fig. 3E**). We tested whether these neurons were indeed inhibitory using cross-correlograms fitted using a generalized linear model (GLMCC) (*37*). We found seventeen (put.) inhibitory influencing PFC to CD neurons and four (put.) excitatory neurons (*α* < .0001 nominal GLMCC threshold test after (*37*), **Fig. S9**), indicating that a majority of neurons in cluster 8 had an inhibitory effect on the CD, potentially by reducing drive from projecting neurons.

**Fig. 3:**
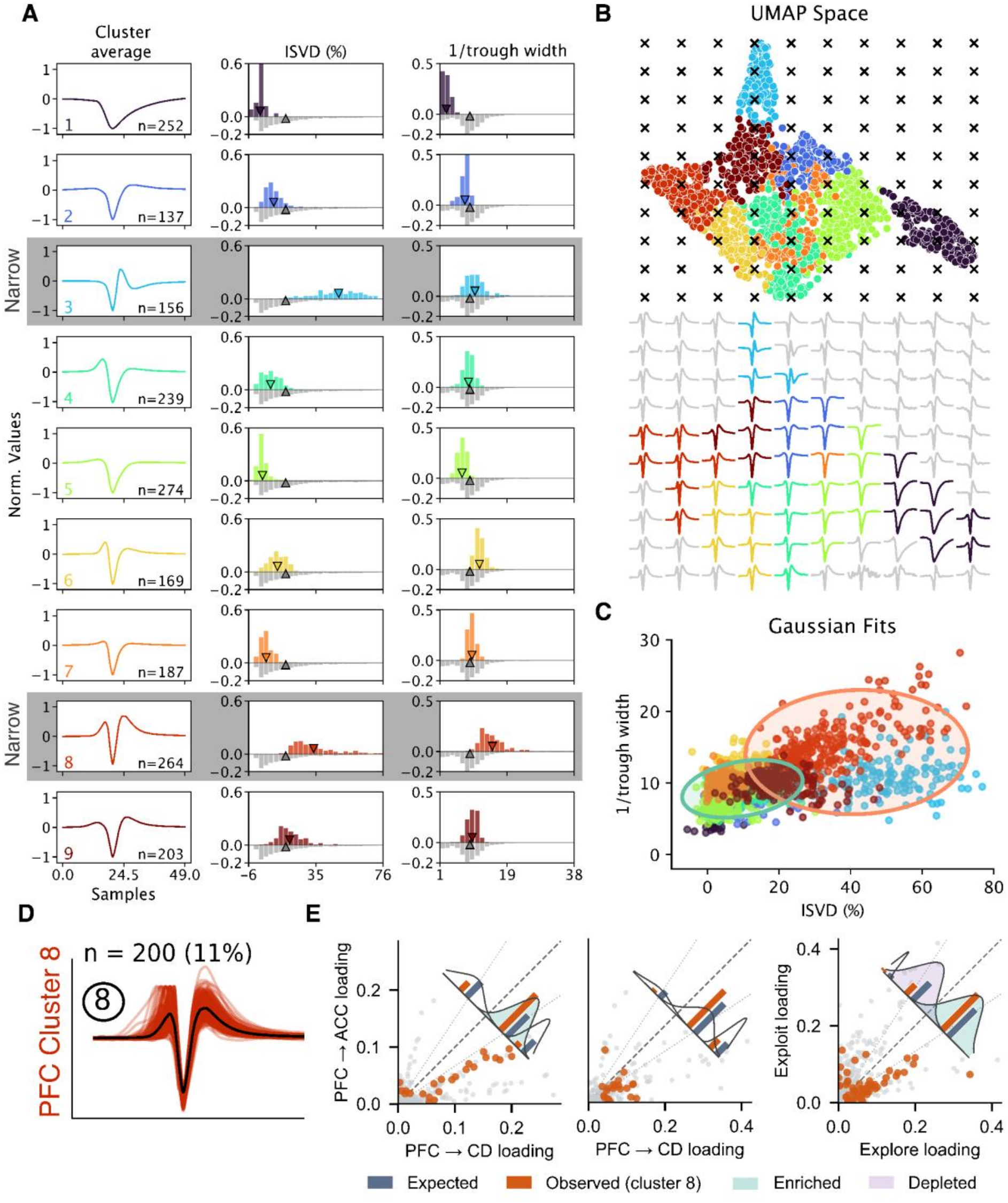
Putative PFC interneurons are more active during learning. (**A**) Average cluster waveform of each WAVEMap cluster for ACC and PFC neurons along with their initial slope to valley decay (ISVD %) and inverse trough width at 50% of trough height. (**B**) Visualization of high-dimensional cluster projected onto the top two UMAP dimensions with inverse transform to visualize waveform shapes. After Lee et al. **(C)** Two Gaussian mixture model fitted to the scatterplot of neurons by ISVD % and inverse trough width. **(D)** Waveforms of cluster 8 in the PFC, identified as putative narrow-spiking waveforms. (**E**) Visualization of omnibus analysis over Gaussians for each cluster showing observed versus expected cluster 8 neuron counts for each Gaussian. The three largest omnibus-scored Gaussians are highlighted.

### High-loading neurons carry the necessary information for object selection

Inter-areal communication subspaces may carry information about the expected value and the location of the fixated object, which would suggest that they could guide gaze towards the most valuable object in space. We tested this by extracting neurons that participated in subspaces with high-loadings (median split) and decoding the value and location of the fixated objects around the time of object selection. The expected value was estimated using a hybrid working memory-reinforcement learning model (**Fig. S10**)(*20*). Decoding revealed above-chance decoding of joint location-value information in the explore state for the PFC and CD, and in all three areas during the exploit state in the high- and low-loading groups of neurons (**Fig. 4A-C**). Decoding was similarly strong for high- and low-loading neuron groups, except for the PFC decoding in the PFC-CD pair after learning, where the high-loading neuron group had better decoding accuracy (**Fig. 4C**). The first time point with decoding accuracies above the 25% baseline level on average preceded fixation onset across the PFC and CD subpopulations of neurons during explore and exploit states (**Fig. 4D**), with more robust early onset of decodability during exploitation in the CD relative to PFC and ACC (holm-adjusted permutation tests, CD-PFC, p < .05; CD-ACC, p< .01) (**Fig. 4E**).

**Fig. 4:**
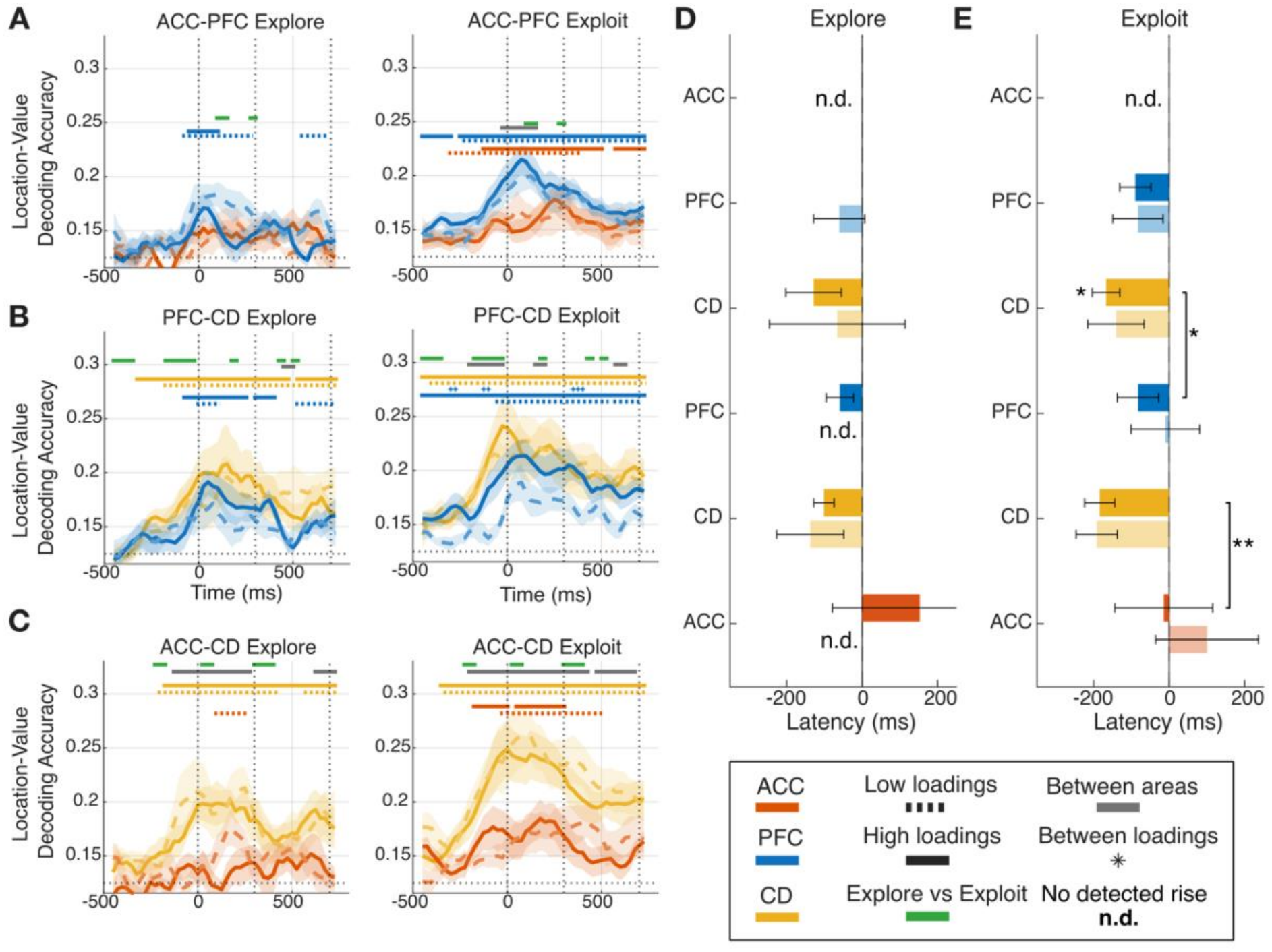
Task-necessary information is decodable from communication subspaces. (**A,B,C**) Decoding accuracy of joint location-value information for each area pair during explore and exploit. Solid significance bars denote decoding accuracy above baseline for high-loading neurons while dotted significance bars indicate decoding accuracy above baseline for low-loading neurons. Green bars indicate significant differences between explore and exploit, grey bars indicate differences between areas, and asterisks indicate differences between high and low loading neurons. (**D**) Latencies with bootstrapped error bars for each area by pair and for high-loading (dark) and low-loading (light) neurons. For the ACC-PFC pair, the ACC did not have a reliable rise to 25% above baseline and is thus not depicted. (**E**) Same as for **D**, but for exploit (significance stars are: ^*^ p < .05, ^**^ p < .01, Holm adjusted permutation tests).

## Discussion

Flexible learning requires exploring multiple hypotheses and revising one’s beliefs after feedback, all internal mental processes. While previous studies have mainly focused on inter-areal communication during externally imposed stimulus presentation within area pairs (*10, 38*), here we tracked how the PFC, ACC, and CD coordinated their spiking activity during self-directed object selection, reflecting internally generated top-down coordination rather than externally imposed coordination. We found that active neuronal communication subspaces arose prior to object fixation and contained the joint location-value information of to-be-selected objects, suggesting that they might participate in shifting where attention is directed and organizing object selection.

This process was not static. Transitioning from exploring to exploiting object values shifted network activity and neuronal participation within the communication subspace. Firstly, the magnitude of inter-areal connectivity within neuronal communication subspaces increased for the ACC-PFC and ACC-CD pairs from exploring to exploiting. This could be due to higher-dimensional cortical representations during learning versus after learning (*39*), or informational content becoming more redundant with learning (*40*). It might also reflect the rapid alignment of a neural ensemble with communication subspace directions, leading to more efficient inter-areal coordination or a combination of these factors. The exception to the changing subspaces with learning was the presence of a persistent PFC-CD subspace in both explore and exploit states. This finding points to a unique subspace that coordinates visual object selection across states during and after learning.

Second, the direction of information flow within the network changed substantially with learning. During exploration, the ACC and PFC led the CD prior to fixation (**Fig. 5A**) with sustained PFC input briefly after fixation (**Fig. 5B**), suggesting increased cortical inputs to the striatum during trial-and-error learning. After learning, the CD had the earliest decoding of joint location-value information and it engaged in feedback with the PFC to the ACC post fixation, underscoring its importance for spatial attention and action selection (**Fig. 5C,D**) (*33, 41*). These results suggest that goal-directed and exploitative behavior both involve coordination across cortico-basal ganglia-thalamocortical loops and recruit subnetworks differentially with more input from cortex during learning and the employment of ‘cognitive shortcuts’ and ‘attentional policies’ during more learned behavior (*42, 43*).

**Fig. 5:**
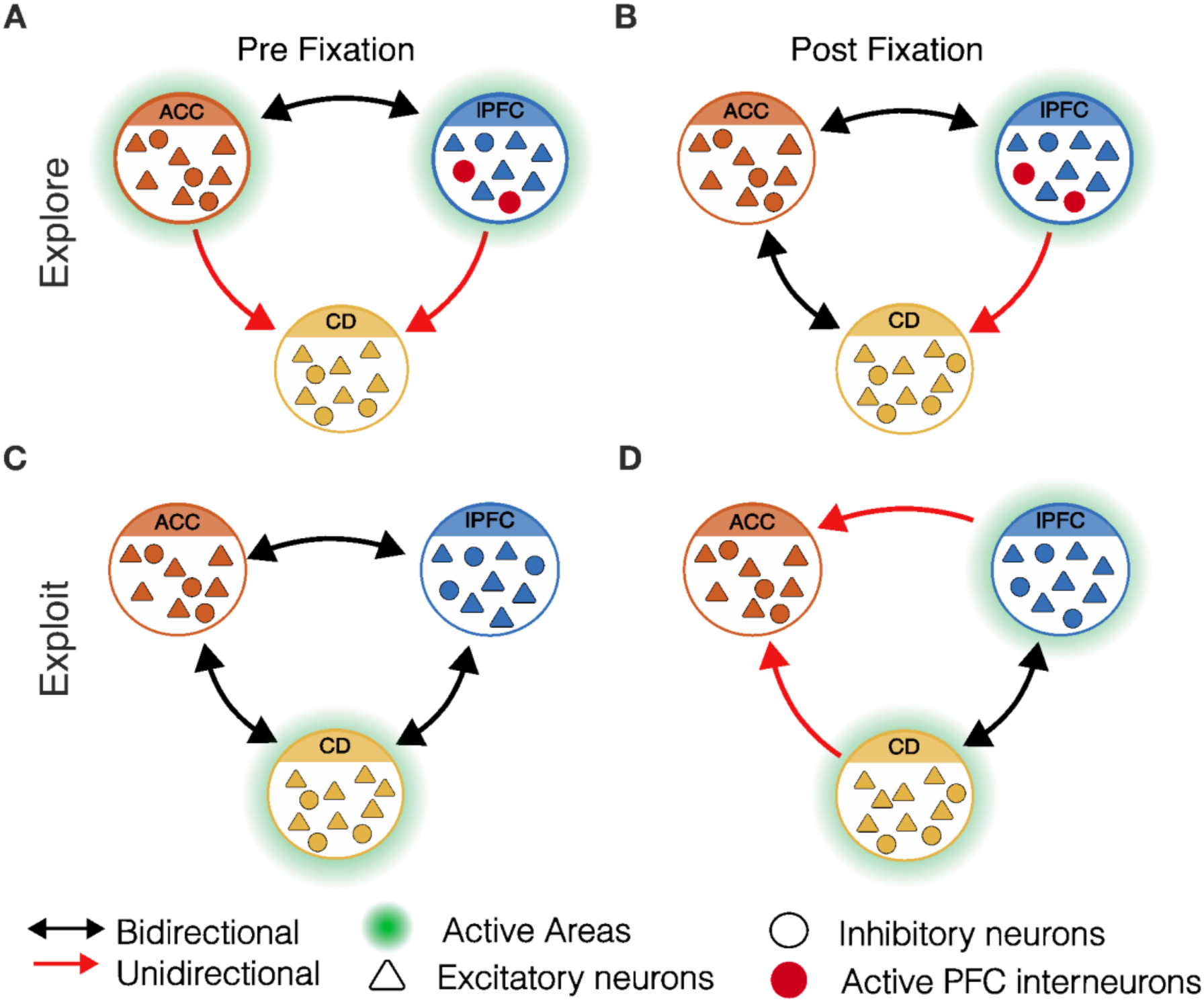
Network connectivity dynamically changes during exploration and exploitation to coordinate object fixation and selection. (**A**) Network diagram of inter-areal communication and neuron participation before fixation. Areas with unidirectional communication are highlighted as active (**Fig. 2B**), and high-loading neuron clusters are highlighted in red (**Fig. 3E**). (**B**) Post-fixation areas with communication dynamics and active neuron clusters. (**C**) Pre-fixation exploit dynamics (**Fig. 2B**) and early CD activity (**Fig. 4E**). (**D**) Unidirectional feedback from lPFC and CD to ACC (**Fig. 2B**) along with elevated decoding from PFC and CD (**Fig. 4C**).

Supporting this view, we also found that distinct neuronal populations underlie exploratory and exploitative behavior (**Fig. 2I,J,K)**. For example, in the ACC and the PFC, more neurons covaried in an ACC-PFC selective subspace during exploration than during exploitation (**Fig. 2C**). This could reflect higher cognitive processes including revising and considering new hypotheses. We also found more PFC (put.) interneurons within the PFC-CD exploration subspace than expected, which suggests they contribute to learning during high uncertainty as found in previous work (*34*) and play an important role in rapidly routing information flow between these areas during learning—a hypothesis suggested by theoretical work (*44*).

Recent articles have highlighted a need for multi-region recordings and the unification of population-based and circuit-based findings (*45*–*47*). Our work accomplishes this by pairing directional dimensionality-reduction methods applied to spiking populations in three areas across the frontostriatal network with single-cell analysis, linking population activity to lower-order circuit mechanisms. Together, these results provide a comprehensive network model of object selection and its dynamic development across learning at multiple spatial and temporal scales, allowing for a thorough understanding of frontostriatal network interactions for self-directed decision making throughout learning.

## Acknowledgements

We would like to thank Robert Louis Treuting for his help with anatomical probe reconstructions. This work was supported by funding from the National Institute of Health (R01MH123687).

## Notes

### Competing Interest Statement

The authors have declared no competing interest.

