## Supplemental Materials for "Neuronal communication subspaces arise across prefrontal-striatal circuits prior to gaze-directed object selection and develop with learning"

**Material and Methods**

Ethics Statement

All animal and experimental procedures were conducted in compliance with the National Institutes of Health Guide for the Care and Use of Laboratory Animals and the Society for Neuroscience Guidelines and Policies were approved by the Vanderbilt University Institutional Animal Care and Use Committee (IACUC).

Experimental Procedure and Design

Two adult male rhesus macaques (Subject W: 14yrs/12.2 kg; Subject I: 15yrs/12.1 kg) performed the experimental task in a sound-proofed chamber. Subjects were head fixed with a 21" LCD screen positioned 63 cm away from their eyes. Then monitor display measured 50.8 x 20.4 cm with a 1920x1080 resolution and ~ Hz refresh rate. At the beginning of each session, gaze was calibrated using 9-points on screen. The task, behavioral monitoring, and reward delivery were controlled by an open-source software: the Multi-Task Suite for Experiments (M-USE) (*1*). Gaze was monitored using a Tobii TX300 infrared eye tracker at a sampling rate of 300 Hz. The flexible learning task consisted of selecting one of three 3D-rendered objects on screen, called quaddles (*2*), which differed across four possible feature dimensions, (body color, body pattern, body shape, and arm shape), two of which were active on a given trial. Each quaddle was positioned randomly at one of the four corners making up a virtual square grid 10.5 cm across with 5° eccentricity relative to the center of the screen. The quaddles were ~2.5-3 cm large which corresponded to ~2.5° degrees of visual angle. Trials started with a blank screen after which the quaddles appeared on screen. Subjects had 5s to explore the options before choosing an object via fixation. Subjects were required to fixate an object (onset of fixation abbreviated Fix) for .7s for a decision to be recorded (DR), however, breaking fixation after .3s aborted the trial and so we also demarcated 300 ms as the time at which the choice was committed (ChC). 200 ms after the decision was recorded, subjects received visual feedback in the form of a halo, yellow for correct and blue for incorrect. Afterwards, if the trial was correct, a circular green token appeared on screen and filled up the token bar, or if incorrect, grey tokens appeared on screen and decremented the token bar based on the current gain and loss condition. The token bars started with three tokens with a total of eight token slots, which when filled, lead to dispensation of fluid reward and a reset to three tokens.

*Block Conditions*

Each session consisted of 36 blocks which lasted for ~30-60 trials. Each block, a singular feature was designated as the rewarded feature (**Fig. 1B**). The rewarded feature changed pseudo-randomly between blocks. In the 'Same' condition, the same objects were shown onscreen, but a new target feature was chosen while in the 'New' condition, all new objects appeared on screen. The shifts could also be intra-dimensional (ID) (i.e. a new target feature within the same feature dimension such as blue body to green body) or extra-dimensional (ED) (the target feature was from a new feature dimension e.g. green body color to checkered body pattern) We focused on the Same versus New conditions as we previously did not find differences in the ED/ID comparison (*3*). Blocks also differed across four possible token reward conditions. Subjects could gain two or three tokens and lose one or three tokens for four different motivational contexts with differing net gains (+2/-3, +3/-3, +2/-1, +3/-1). Motivational contexts were equally assigned to each Same/New and ED/ID transition but were pseudo-randomly assigned within each condition.

*Electrophysiological Recordings*

We analyzed electrophysiological recordings from twenty-nine session across subject I and W. In these sessions, neural activity was recorded with two 256-channel linear silicon probes inserted into each area (Diagnostic Biochip, 25 µm interelectrode distance) using Open-Ephys62 (open-ephys.org) and the Intan Recording System (RHD2000, Intan Technologies, LLC). Neuronal wideband recordings were notch filtered (60, 120, and 180 Hz) with a filter bandwidth of 2 Hz to remove power noise and its harmonics and re-referenced using median referencing across channels.

*Probe Reconstructions*

Recording sites across the ACC, PFC, and CD were reconstructed in 3D Slicer 63 (http://www.slicer.org, version 4.11) (**Fig. S1**). For each subject, MRI scans acquired at the time of recording chamber implantation were co-registered with CT scans acquired after implantation. The recording chamber was then segmented, and the image space was reformatted relative to the chamber. We recorded the x, y, and z coordinates of each stimulation electrode tip relative to the chamber, with z representing electrode depth, to reconstruct each electrode's penetration depth. The reconstructed electrode tip locations were marked on the MRI images, and monkey W's stimulation sites were overlaid onto monkey I's image to provide a unified view of the stimulation sites.

### Behavioral Analysis and Modelling

*Learning Curves*

Learning curves were calculated by fitting a logistic sigmoid to the average performance across blocks for each trial, averaged across sessions. The trial-to-criterion was calculated as the first trial where the monkeys to reached 80% accuracy within a ten-trial prospective window. For this analysis, we excluded any block where the monkey guessed the first trial correctly as this did not reflect trial-and-error learning.

*Parallel Belief State Model of Attention*

We used the recently proposed Parallel Belief State (PBS) model of attention (Tiesinga and Womelsdorf, 2025) to objectively quantify which object features are considered by subjects for an exploratory or an exploitative eye movement and choice. The PBS model is an input-dependent Hidden Markov Model (*4–6*) that infers hidden (belief) states, which reflect the belief for each given feature about whether it is the target feature, and updates that belief through a state transition during the current trial which is a function of the choice and outcome on the preceding trial (referred to as the input, $h$). The mechanics of the model are described in (*7*)**.** In brief, the model accounts for observations of whether a feature was present or not in a chosen object by maximizing the probability for the entire set of observed choices ($c_{1:T}$) across trial 1 to T, and the corresponding hidden states ($z_{1:T}$) given the inputs ($h_{1:T}$) and a particular model m with parameters $\theta_m$, which is written as: $p(c_{1:T}, z_{1:T}|h_{1:T}, \theta_m)$ The notation is shorthand for a collection of parameters including the state transition and emission matrices as well as the distribution of initial states, which are detailed below. The choice (value 1 if the feature is chosen, that means it was present in the chosen object, and likewise 2 if it is not chosen) on trial t is predicted based on the current input $h$ which reflects the choice and outcome (one when rewarded and zero otherwise) on the preceding trial encoded as a number between 1 and 4. This input directly influences the predicted choice via the input-dependent emission matrix (size: number of choices (2) times the number of states $n_s$), which links the current hidden state which reflects the longer-term history to a probability of the choice via the matrix element of this matrix. The input dependent transition matrix (size $n_s$ x $n_s$) updates the hidden state to reflect the most recent available choice/outcome, which reflects the learning dynamics of the subject. This new hidden state is not directly accessible for the observer, instead it must be inferred, which yields a probability for it to be in state j when considering both the past and the future.

To optimize the prediction success of the model we maximize the likelihood of the actual choice via an EM procedure. We used the log likelihood (LL) of the entire sequence of choices, normalized by the number of choices, its value should exceed that of chance (note that we use a natural e-based log). For each of these data sets we ran the procedure 10 times for different initial values for the parameters (the emission and transition matrices, and the prior on the distribution of the hidden states), to avoid the EM procedure getting stuck in local maxima and performed a minimum of 300 iterations for our analysis.

The parallel belief states comprise the combined states of each of the 12 object features (see Suppl. Text). We assign a label to each PBS of each trial based on the distribution of states across features as follows:

- **Exploit target**: target feature is in “exploit” state
- **Explore target**: target feature is “explore” state, no other features are in the “explore” state

- **Explore distractor**, one or more non-target features states are in an "explore" state, the target feature is not in an "exploit" state
- **Perseverance**, feature corresponding to target of preceding block is in "persist" or "exploit" state
- **Persist in distractor feature**, a different feature (neither current target, not previous target) is in an "exploit" state
- **Other**: none of the above

The *Other* state category occurred too rarely to meaningfully separate it further. For our analyses, we combined explore target and explore distractor into the explore condition and we focused on analyzing the differences between explore and exploit states.

<u>Neural Data Analysis</u>

*Canonical correlation analysis*

Spiking data were residualized by z-scoring and subtracting the peri-stimulus time histogram across time (separately by condition when applicable). By residualizing, we were able to focus on co-fluctuations in neural activity that were not explainable due to average trial dynamics (*8*). We performed both static (zero time-lag) and time-lagged canonical correlation analysis (CCA). For the static CCA, we used a 125 ms sliding window that was slid 5 ms at a time and that contained non-overlapping 25 ms bins of residualized spiking data across trials. These parameters ensured that we had a high level of resolution to capture rapid changes in inter-areal coupling. Given areas X and Y with *n* and *m* neurons, *z* trials, and *k* bins, we can define the sliding window data at timepoint *t* as:

$$\mathrm{X_t} \in \mathbb{R}^{kz \times \mathrm{n}}, \;\; Y_t \in \mathbb{R}^{kz \times m}$$

CCA removes within-area correlations by calculating each area's whitened within-area covariance matrix and then performing singular value decomposition on the cross-covariance matrix multiplied by the whitened matrices. Given mean-centered covariance matrices with N = *kz* observations:

$$C_{XX} = \frac{X_t^\top X_t}{N-1}, \quad C_{YY} = \frac{Y_t^\top Y_t}{N-1}, \quad C_{XY} = \frac{X_t^\top Y_t}{N-1}.$$

The whitened cross-covariance matrix T is given by:

$$T = C_{XX}^{-1/2}\, C_{XY}\, C_{YY}^{-1/2}$$

And eigen decomposition yields:

$$T = USV^\top$$

The max correlation value is given by the singular value of the first canonical components, $\rho_1 = \sigma_{max}(T)$. For the time-lagged version, we used absolute time lags of $\pm$ 80 ms with steps of 5 ms between areas and computed $\rho_1$ for each time lag. For a given time-lag $\tau$:

$$T = C_{XX}^{-1/2}(t)\, C_{XY}\,(t, \tau) C_{YY}^{-1/2}(t + \tau)$$

$$\rho_1(t, \tau) = \sigma_{max}\big(T(t, \tau)\big)$$

Not all sessions showed a peak in correlation around fixation. We determined which sessions to include for analysis via exploratory static and time-lagged CCA. Static and time-lagged CCA was performed for each area-pair and session, and we included sessions where we saw a peak in correlation in the $\pm$ 200ms around fixation for both the static CCA and the averaged time courses of the lagged CCA. Additionally, sessions were excluded if they exhibited pronounced regularly repeating oscillations throughout the entire time course suggestive of recording artifacts. We found peaks in 11/16 sessions for the ACC-PFC pair, 12/17 sessions for the PFC-CD pair and 12/28 sessions for the ACC-CD pair pooling both monkeys' sessions together.

*Latency*

To calculate the onset latency of inter-areal connectivity, we performed a latency analysis of the time-lagged average time-courses for explore and exploit. Latency was defined as the earliest time-point that the session-average time course exceeded 25% of peak – baseline and remained above that threshold for the subsequent 100ms. Baseline was defined as the first 380ms of the trial, (800-420ms before fixation). The peak was the maximum CCA $\rho$ in the analysis window of -500 to 700ms around fixation. 95% confidence intervals were estimated by 10,000 bootstrap resamples of sessions. One-sided sign-flip tests tested whether individual-session latencies were earlier than fixation. Two-sided unpaired permutation tests compared latencies between pairs of areas. Two-sided paired permutation tests were also used to compare latencies between Explore and Exploit, swapping state labels within sessions. Holm correction across area pairs was applied separately to tests within each state before fixation, across pairs of areas within each state, and across pairs of areas for Explore-Exploit comparisons.

*Neuron participation analysis*

To understand how individual neurons participated to the communication subspaces, we extracted the loadings of the neurons onto the subspaces which is defined as the Pearson correlation between the resdualized neural activity of each neuron with the canonical component of its own area for each time window. Formally, let $X^A$ and $X^B$ denote the residualized activity matrices for areas $A$ and $B$, with rows corresponding to observations pooled across time bins and trials and columns corresponding to neurons. The vectors $w_1^{A|B}$ and $w_1^{B|A}$ denote the first canonical weight vectors obtained from CCA between these areas, such that $u_1 = X^A w_1^{A|B}$ and $v_1 = X^B w_1^{B|A}$ are their respective canonical variates. Each neuron's loading is written as $\ell_i^{\{A|B\}}$ or $\ell_i^{\{A|B\}}$, with, *i* and *j* indexing neurons within their respective areas, and (A|B) denoting area *A*'s participation in the CCA fitted with area *B*, without implying directionality.

$$\ell_i^{(A|B)} = \mathrm{corr}(X_i^A, u_1) \qquad \ell_j^{(B|A)} = corr(X_j^B, v_1)$$

The loadings gave us a quantitative measure of how strongly each neuron's activity covaried with the canonical component across time. We interpreted the loadings rather than the canonical weights because weights can suffer from multicollinearity when there are multiple correlated within area neurons. We also used loading magnitudes because the signed pairs of the canonical components are jointly arbitrary, (*-u,-v*) and (*u,v*) yield the same canonical correlation.

For the analysis of inter-areal loading selectivity, sessions that had inter-areal peaks for both area pairs were included. Since CCA was fit separately for each area pair, the canonical component for area *A* with area *B* is different than the canonical component for area *A* with area *C*. Thus, we were able to plot the loading magnitudes of each area's neurons with both other areas. Data of loadings from a window of 100ms before fixation to 150ms after fixation was used for the analysis. To quantify selectivity, we used the method traditionally used for interocular selectivity (*9*). Where $\bar{\ell}_A^i$ is the mean loading of a given neuron for area *A* and $\bar{\ell}_B^i$ is the loading of that same neuron with area *B,* the selectivity index (SI) is:

$$SI = \frac{|\bar{\ell}_A^i| - |\bar{\ell}_B^i|}{|\bar{\ell}_A^i| + |\bar{\ell}_B^i|}$$

In addition to the selectivity index, we tested if the data were normally distributed with a Lilliefors test of normality finding that all the data were non-normally distributed (Lilliefors $p < .01$). To model the underlying data distributions, we fitted one, two, and three Gaussian models to the data. In all cases, three Gaussians best fit the data as quantified by both a lower Bayesian information criterion (BIC), and Akaike information criterion (AIC) in all cases. The same procedure was applied to measure loadings in explore versus exploit states by plotting each neuron's loading over the same -100 to +150 ms window around fixation for explore states and exploit states separately and for each area combination.

In addition to the time-lagged CCA analysis, the transition point between exploration and exploitation was analyzed using a sliding-trial window. A four-trial sliding window starting six trials before the transition and ending six trials after was stepped forward one trial at a time and average time-lagged time courses were calculated across blocks with the four-trial window. Peak heights and latencies were identified using the maximum value return by the findpeaks function in MATLAB within the -200 to +350ms around fixation.

#### *Neural Data Time-Series Statistics*

All CCA $\rho$ timecourses were baseline z-scored with a baseline of 350ms at the start of the trial segment in the case of static CCA, or with 380ms from the start in the case of time-lagged CCA due to fewer CCA windows at the edge of the trial where the negative time-lags were out of bounds. For time-course data, we used nonparametric paired-session cluster permutation tests across time to evaluate statistical significance (*10*). We tested whether CCA $\rho$ was above baseline by calculating t-statistics across time and evaluating them against baseline statistics. T-stats more extreme than $p < 0.1$ were grouped into clusters. Then each cluster's mass was tested against the maximum clusters generated from the null distribution with $p < 0.05$. Null distributions were formed based on clusters from the baseline period. For between condition tests, we swapped explore and exploit labels with $2^N$ permutations where N was the number of sessions. T-values that exceeded either $p < 0.05$ tail of the distribution was aggregated into clusters and the final cluster mass was compared against the baseline distribution maximum clusters at $p < 0.05$. Statistics were done on the raw data, not smoothed data. Data were smoothed for visualization using a six-window Gaussian kernel.

### Neuronal cell-type analysis

#### *Spike Sorting*

Electrophysiological data traces were sorted into individual spiking neurons using Kilosort 4 (*11*). After Kilosorting, neuron waveform clusters were further manually inspected in the Kilosort GUI: Phy. In Phy, neuron clusters were merged if they were similar in shape, location across electrodes, had similar auto correlograms and merging them did not result in refractory period violations. From the default Kilosort outputs of good or MUA (multi-unit activity), we manually inspected and ranked 'good' neurons as we found that many of these neurons did not pass more rigorous quality inspection in Phy. Putative neurons clusters were ranked from 0-5 with 0 representing pure noise, 1 noisy MUA, 2 good MUA, 3 excellent MUA, and 4 and 5 representing well-isolated single units. These distinctions were made based on a variety of metrics within Phy. We considered the amplitude of clusters relative to the noise floor, how clean the refractory period within the auto correlogram was, to what extent the PCA components of separate neurons were distinct from each other, whether waveforms exhibited line noise and if waveforms drifted or shifted over time and if this was in a 'bow' drift or diagonal drift. Units labelled 4 and 5 were clearly above the noise floor, had no major refractory violations, had distinct PCs from all other units, and had clean mean waveforms. We only included 4 and 5 ranked units in our analysis.

*Classifying neuron types*

To classify neurons into putative cell types, we used WaveMAP, a non-linear dimensionality reducing clustering method (*12*). WaveMAP uses uniform manifold approximation and projection (U-MAP) to great an asymmetric high-dimensional directionally weighted graph. It then uses Louvain community detection to group the graph into clusters which minimize within-area edge distance while maximizing the between-cluster distance. We found that the default Louvain community detection value of 1.0 maximized the modularity of clusters and used it for our analyses. For our U-MAP parameters, we chose the number of neighbors to be 150 to focus on global structure given our large dataset size (N = 1881 cortical neurons and N = 2759 caudate neurons). We fit WaveMAP onto our cortical and CD neurons separately as the striatum only contains inhibitory neurons. For both fits, WaveMAP yielded nine separate neuron clusters.

After applying WaveMAP, we sought to further distinguish the broad-spiking excitatory neuron classes from narrow-spiking inhibitory interneurons. To do this, we calculated the initial slope to valley decay (ISVD) which measures the extent to which the voltage drops 0.26ms after the first post-trough peak which has been previously shown to identify narrow spiking interneurons (*13*).

$$\mathrm{ISVD} = 100 \times \frac{V_{\mathrm{peak}} - V_{\mathrm{peak}+0.26\,ms}}{V_{\mathrm{peak}} - V_{\mathrm{trough}}}$$

We also measured how narrow spikes were by measuring the inverse trough width which was derived from the time in ms between each side of the waveform trough at half of the max trough depth. The max depth was defined as the most negative voltage of the waveform recorded while the time-point at which the 50% threshold was met was calculated using shape-preserving piecewise cubic interpolation (PCHIP) to attain a precise half-max time for the descending and rising portions of the waveform trough respectively.

Neurons were grouped using Gaussian mixture modeling of the ISVD and inverse trough widths. Features were standardized to zero mean and unit variance and reduced to a single dimension using the first principal component. Gaussian mixture models containing one to three components were fitted with 10 initializations per fit. The number of components was selected by minimizing the mean Bayesian information criterion across five cross-validation folds, and the selected model was refitted to all neurons. Each neuron was assigned to the component with the highest posterior probability. Neurons without valid ISVD or inverse trough width values were excluded from the analysis. Our analysis determined that a two Gaussian mixture model fit the data best for both the cortical and CD neuron groups. The two gaussians distinguished broad from narrow spiking neurons, with one gaussian grouping together clusters that had low ISVD and inverse trough width neurons—characteristic of broad rather than narrow waveforms—and another gaussian that captured two clusters of neurons with much larger ISVD and inverse trough, characteristic of narrow spiking neurons (**Fig. 3C, Fig. S3C)**. 22.7% of the cortical neurons and 17.5% of CD neurons were identified as narrow which is consistent with the overall proportions found in other studies (*14*).

*Identification of inhibitory versus excitatory influence*

Influence between PFC narrow spiking neurons and CD neurons was identified using GLMCC, a technique that fits a generalized linear model onto cross-correlograms and compares this to a baseline fit to find significant interactions between neuron pairs (*15*). The GLMCC model was fit onto trial segmented data. We ensured that no spikes coincidences were made across trial boundaries. The original GLMCC parameters were used for the fitting with a smoothly varying baseline and exponential coupling kernels in both directions. The cross-correlograms were evaluated over $\pm$ 50 ms and significant connections were determined based on a nominal two-sided GLMCC criterion with $\alpha$=0.001 as in the original paper.

Decoding

Decoding was performed using time-resolved linear support vector machine (SVM) decoding with one-versus-one error-correcting outputs (MATLAB *fitecoc*) for multiclass classification. All classes were balanced before being split into an 80-20 test train split. During training, five-fold cross validation was used to determine the optimal regularization parameter C out of possible values of {0.1,1,10} at each time window. Each time window consisted of five 125 ms bins of residualized neural activity. The optimal model for each window was then refitted onto the training data and evaluated on the held-out trials.

Decoding was done on four variables, object location, value, choice probability, and joint location times value. The object location corresponded to each quadrant of the screen which the quaddles could appear in. The value was derived from our reinforcement model's value estimates of each feature, and we took the max valued feature on each object as the object value for decoding. Choice probability was also derived from the reinforcement learning model and was unique for each object. Finally, joint location-value was a combination of median split value with location yielding 8 possible location-value combinations. Decoding was done separately for explore and exploit trials and value and choice probability were split into high-low via a median split based on the proportions within explore and exploit separately as well.

*Latency*

To estimate the latency of information encoding within each area, we estimated the timepoint at which 125% of baseline accuracy was reached. Decoding accuracy curves were lightly smoothed using a four-bin moving average (100 ms). Baseline accuracy was calculated over the first 10 bins of the time course. Latency was defined as the first of five consecutive 25-ms bins after the baseline period in which session-averaged accuracy reached at least 125% of baseline. Latency estimates were summarized as the mean ± SD of 10,000 session bootstrap resamples, provided that more than half yielded valid latencies. For high-loading populations, one-sided sign-flip tests assessed whether mean individual-session latencies preceded fixation, with Holm correction across all pair-specific high-loading populations separately within Explore and Exploit states. We tested whether high-loading CD neurons encoded location-value information prior to high-loading ACC and PFC neurons by using one-sided paired permutation tests of earlier CD latency with both areas. Complete accuracy curves were swapped within matched sessions, and group-average latencies were recalculated for 10,000 permutations. Permutations yielding undefined latencies were excluded.

*Decoding Time-Series Statistics*

For decoding time-series data, we used nonparametric cluster permutation tests across time. Above-chance decoding was assessed by subtracting chance accuracy (1/K, where K was the number of classes) from each session's accuracy and calculating one-sample t-statistics across sessions. Consecutive bins exceeding a one-sided cluster-forming threshold of $p < 0.10$ were grouped into clusters, with cluster mass defined as the sum of their t-statistics. Null distributions were generated by sign-flipping each session's entire chance-subtracted accuracy curve over 5,000 permutations, retaining the maximum cluster mass from each permutation. Observed clusters were evaluated at $p < 0.05$.

Differences between loading groups, areas, and behavioral states were assessed using paired t-statistics, with positive and negative clusters formed separately at a two-sided threshold of $p < 0.05$. Null distributions were generated by randomly swapping the two labels within each paired input column over min ($2^N$, 1,000) permutations, where N was the number of paired columns. Each permutation contributed its largest positive and most negative cluster masses to separate null distributions. Clusters were evaluated at $p < 0.05$ per tail. All permutations used the full input time course. Statistical tests were performed on unsmoothed accuracy curves; a four-bin moving average (100 ms) was applied for visualization.

### Supplemental Text

*Preprocessing the dataset for hidden Markov modeling*

We fit the model to choices of the subjects in trials across multiple blocks. To preprocess the data for the model optimization we translated the chosen object and outcome of each trial, for each considered feature, into when the chosen object contained the feature, and when it did not. We performed two types of fits, one focused on how a single target feature is learned, the second on how feature belief states change in parallel, whose results are reported in the main text.

For the first fit, we consider each feature separately and find all the blocks in which it was the target, and for each target block include the preceding block, which are then all concatenated. We then concatenate these single feature data sets across all features. The primary reason for doing this is that the feature under consideration has an equal likelihood to be a target or not, so that the model prediction must be better than a random choice with probability half for each option (hence corresponding to a log-likelihood). Furthermore, the ‘exploit’ state, indicating that the feature is chosen persistently because it is the target feature, should occur in a reasonable fraction of trials so that its properties can be well estimated.

In each block there were six features present in the objects, when in a subsequent block one or two new dimensions are introduced (in the condition ‘New’), there are either three or six new features added. For the second fit, we organized the data so that we could track these block transitions, while avoiding having to track the states of too many features simultaneously resulting in too few samples for adequate statistics, hence we cut the trials into groups of consecutive blocks in which no more than 12 unique features are present. This means that the set of features (and their index) will vary across these groups. The choices for each of the (at most 12) different features are combined in one big set and fitted by a common transition matrix. This way of combining choices means that the “exploit” state will occur less often for each feature, because it is the target in a small fraction of blocks. This makes fitting additional states reflecting the learning process of target challenging.

*Fitting the parallel belief state model*

We set out to find the best set of parameters for the model, in terms of the dimensionality of the hidden state, the depth of the history (how many previous trials are incorporated, in our standard formulation this was 1) and the best model type (the input dependence of transition and emission matrices). We were guided by our previous analyses (*7*), hence we only explored 3 and 4 states and considered only the case in which the emission matrix was not input dependent. In the first fit there was a difference in number of feature choices across subjects (M1: 71074, M2: 28835), which lead for M1 four states to have a lower (better) negative log likelihood (NLL) and BIC than for three states (NLL = 0.4131 vs 0.4189, BIC = 59336 vs 59866, for three states there were 29 and for four states there were 55 parameters). However, for M2 the NLL was still lower for four states, but the BIC was higher (NLL = 0.3914 vs 0.3958, BIC = 23139 vs 23121), indicating that improvement in fit was not sufficient to compensate for the increased number of parameters.

For the second fit, we again compared 3 and 4 states, but fixed the emision matrix, based on our prior experience (Tiesinga and Womelsdorf, 2025), hence the number of parameters reduced to 26 and 51, respectively. The number of feature choices available for M1 and M2 was 425016 and 212256, respectively. For this case, four states led to a lower NLL and BIC for both subjects (M1: NLL = 0.2860 vs 0.2886, BIC = 243864 vs 245697; M2: NLL = 0.2747 vs 0.2784, BIC = 117281 vs 118560). However, when we inspected the fraction of trials the feature was chosen in each state, it turns out the “exploit” state was artificially split into two states, with approximately the same fraction of choices and so we combined these states.

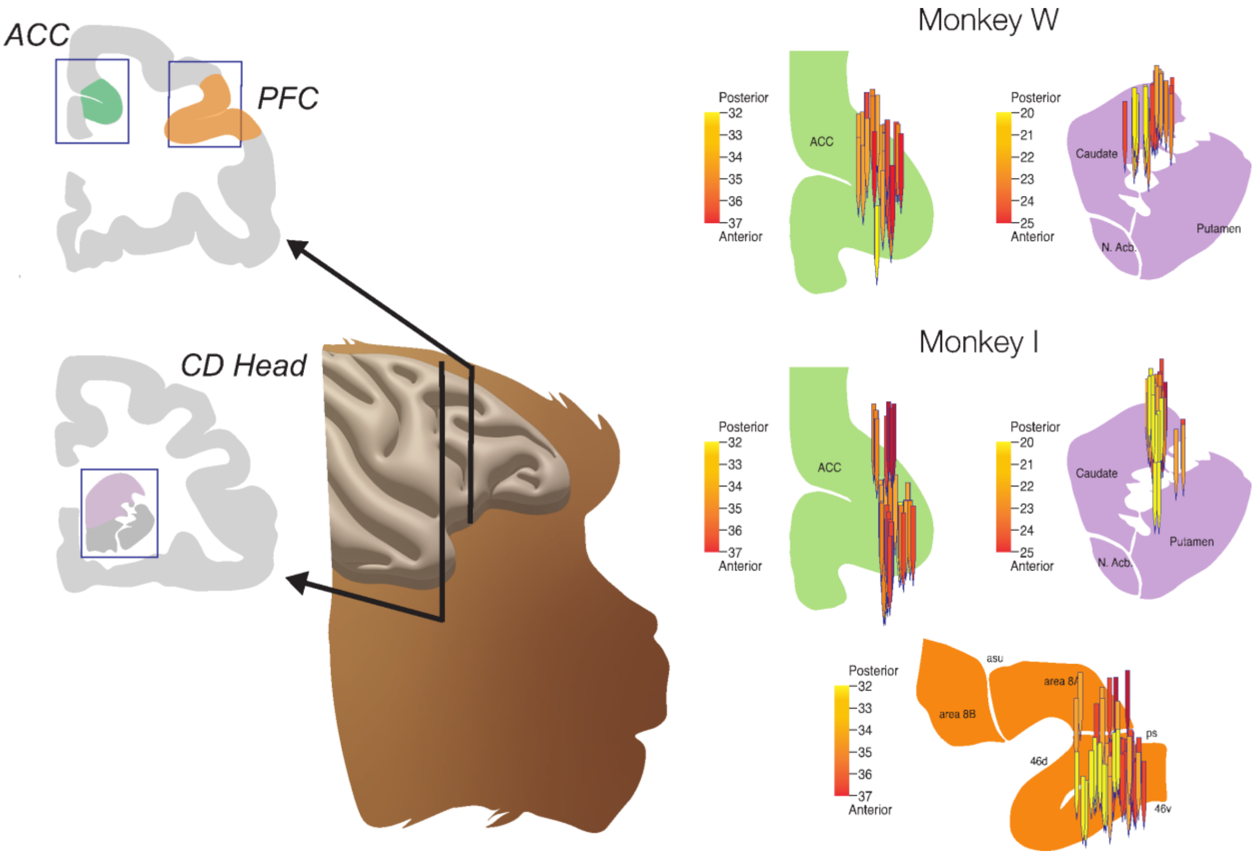


**Fig. S1** The left panel schematic shows the recording locations for the anterior cingulate cortex (ACC), Brodmann area 9/46 of the prefrontal cortex (PFC) and the head of the caudate (CD). To the right are probe reconstructions of all sessions for monkey W and monkey I. Two probes were inserted into each brain area per session, for monkey I recordings were done in all three areas whereas for monkey W, recordings were done only in the ACC and CD.

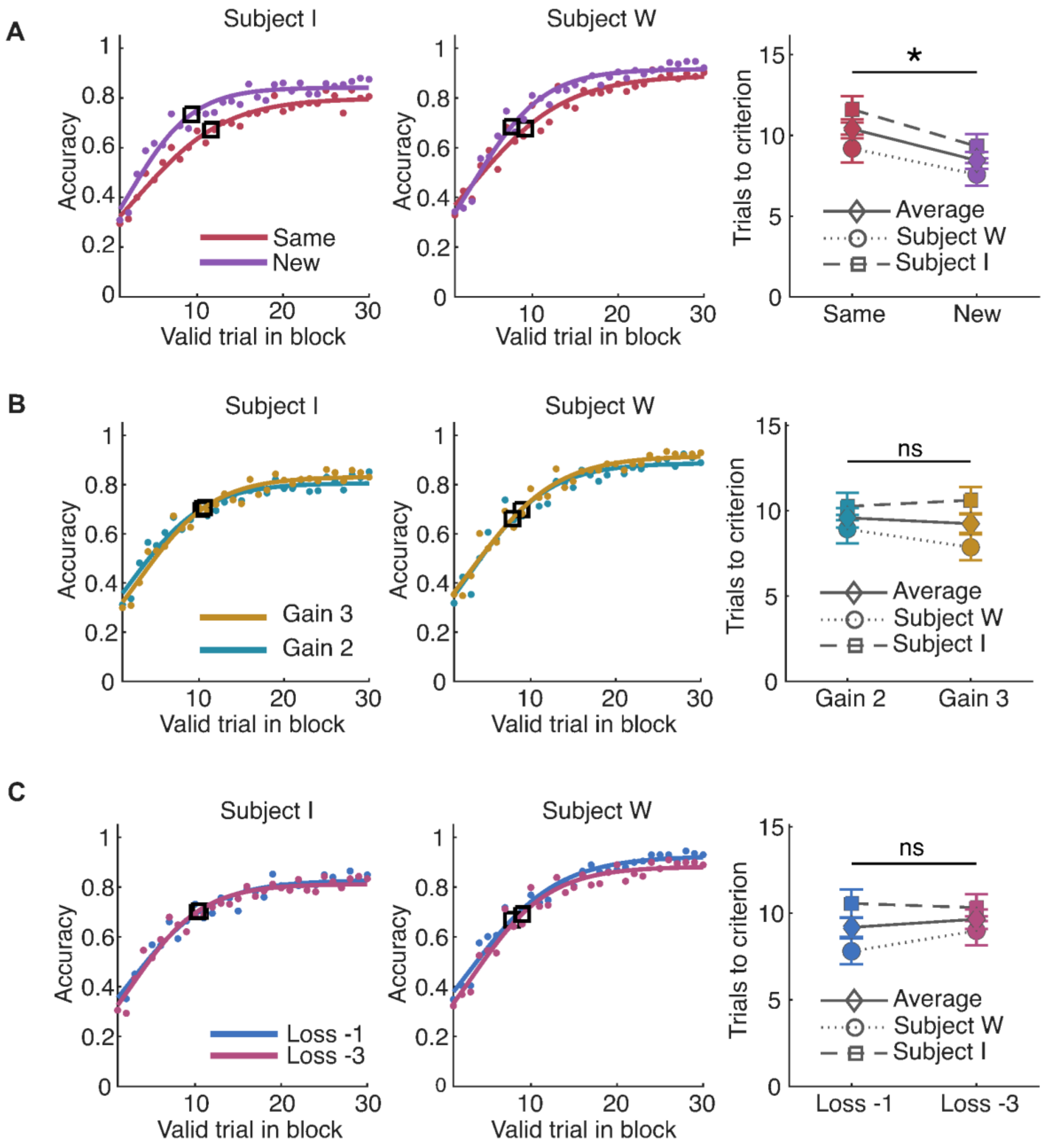


**Fig. S2 (A)** The left two panels are learning curves with logistical sigmoid curves fit to the data and the mean trials to criterion outlined in black for subject I and subject W respectively in the same versus new condition. The rightmost panel is the mean and s.e.m. for each subject individually and on average (* one-sided Holm adjusted t-test with Welch-Satterwaite degrees of freedom p = .0362). (**B**) Learning curves for +2 or +3 token gain and two-sided t-test for a difference in learning rate across both subjects. (**C**) Same as above, but for different loss conditions.

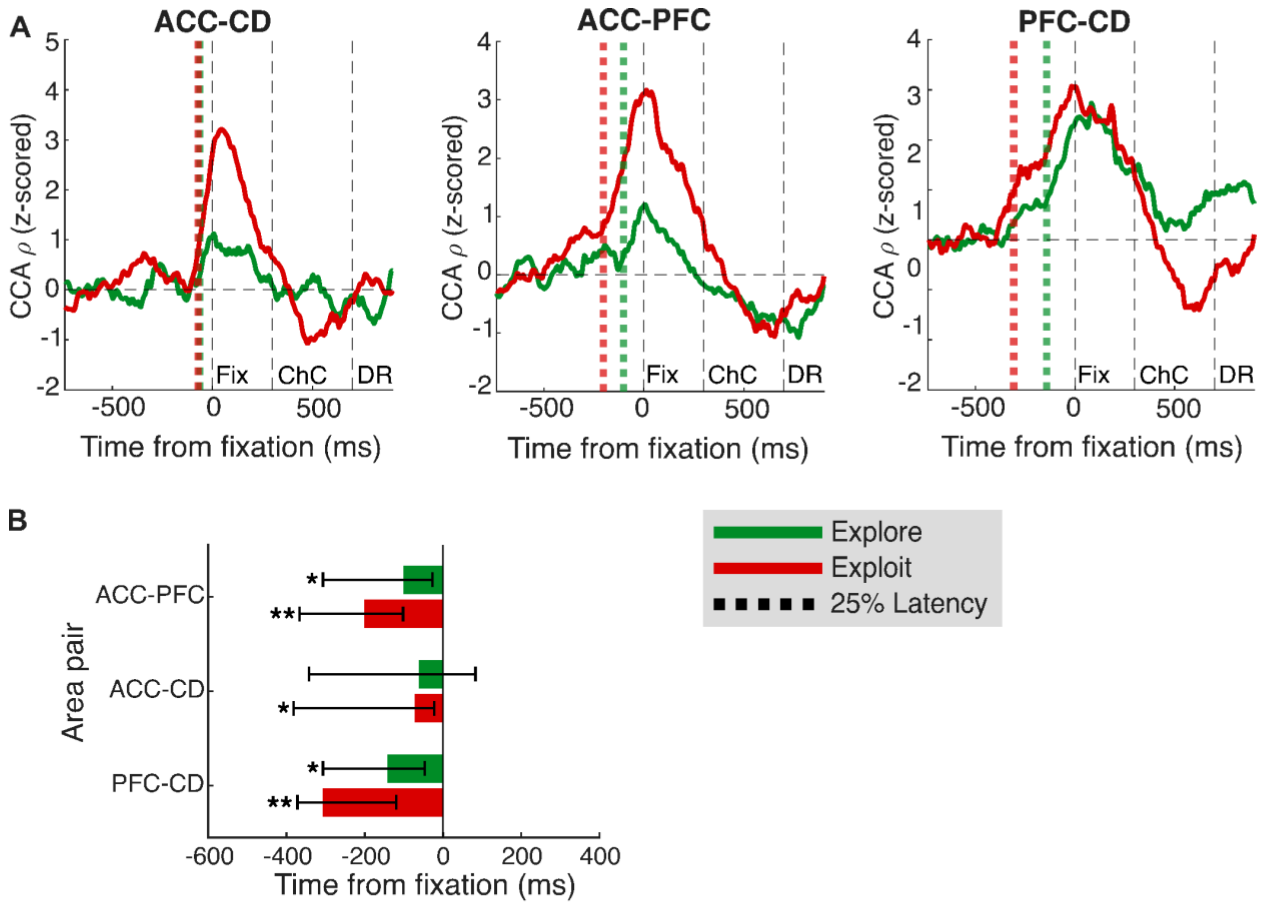


**Fig. S3** (**A**) Averaged explore-exploit z-scored CCA $\rho$ magnitude with the average time to reach 25% of baseline to peak magnitude denoted by dotted lines for explore in green and exploit in black across all area pairs. The average latencies of the inter-areal connectivity increases were -62ms, -102ms, -142ms during explore, and -72ms,-202ms,-307ms during exploit for ACC-CD, ACC-PFC, and PFC-CD, respectively. (**B**) Mean latencies to 25% above baseline with bootstrapped confidence intervals derived by resampling sessions with replacement. Latencies were tested against zero using a one-sided sign flip test and holm-adjusted p-values (* p adjusted < .05, ** p adjusted < .01). All latencies for both explore and exploit were found to be pre-fixation except for the ACC-CD in explore.

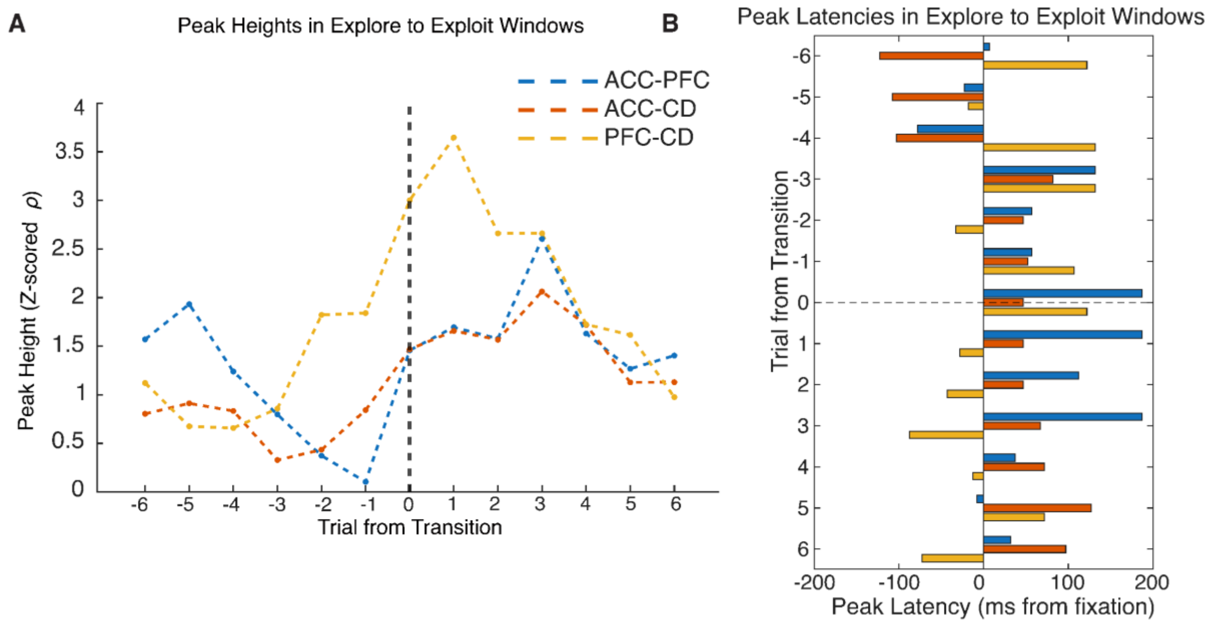


**Fig. S4** (**A**) Peak heights from 4-trial sliding window CCA analysis around transitions from explore to exploit. We a gradual increase in ACC-CD peak heights (shuffle permutation test p = .002) from explore to exploit along with a peak in peak heights around the transition for the PFC-CD pair (permutation test p = .008). (**B**) Latencies of peaks for each sliding trial window relative to fixation. We found that ACC-PFC peaks were later in time around the transition from explore to exploit (permutation test p = .033) and that PFC-CD peaks were later in explore than exploit (permutation test p = .026).

**A**

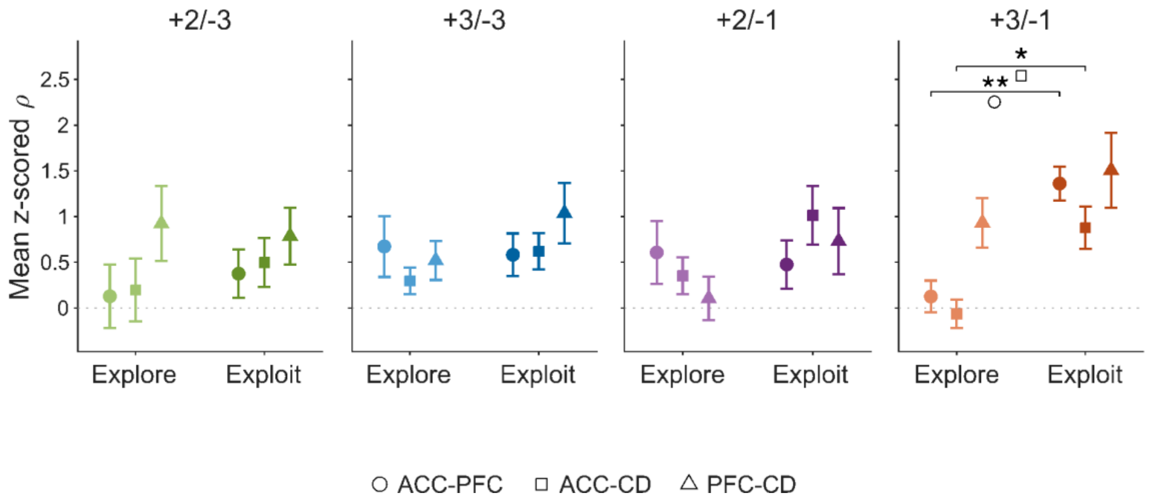


**Fig. S5** (**A**) CCA $\rho$ magnitude in the $\pm$100 ms around fixation across different reward conditions and between explore exploit. For the highest gain condition, we see an increase in the magnitude of peaks for ACC-PFC and ACC-CD with learning as in the overall data (Holm-adjusted two-sided permutation test p = 0.003 for ACC-PFC, p = 0.033 for ACC-CD).

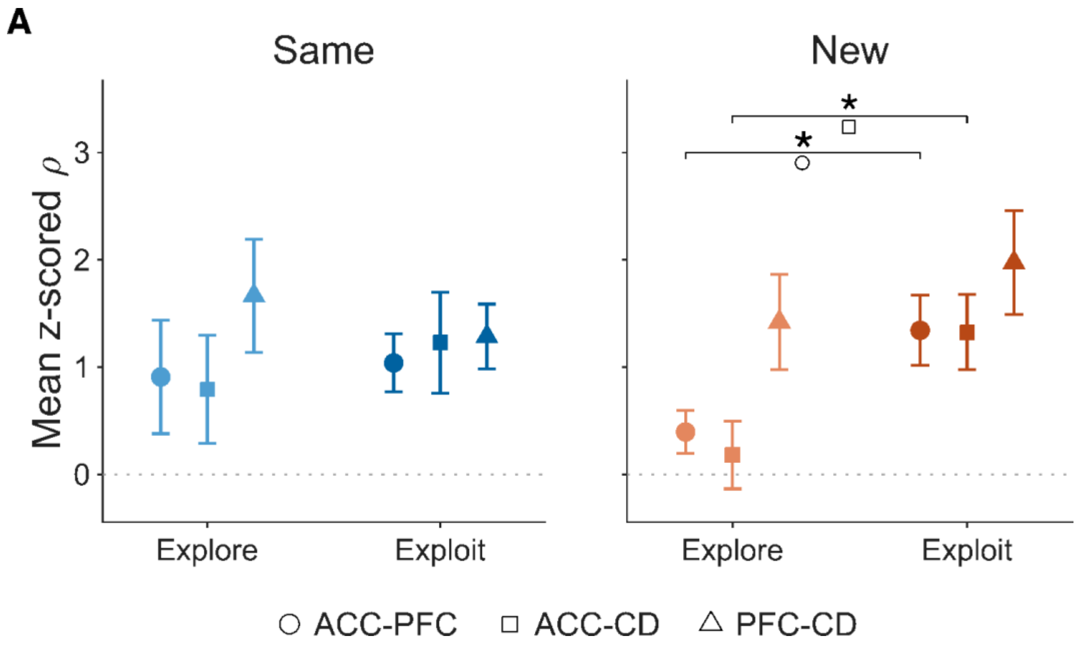


**Fig. S6** (**A**) CCA $\rho$ magnitude in the $\pm$100 ms around fixation across for same and new between explore exploit. For the new condition, we see an increase in the magnitude of peaks for ACC-PFC and ACC-CD with learning as in the overall data (Holm-adjusted two-sided permutation test p = 0.048 for ACC-PFC, p = 0.028 for ACC-CD).

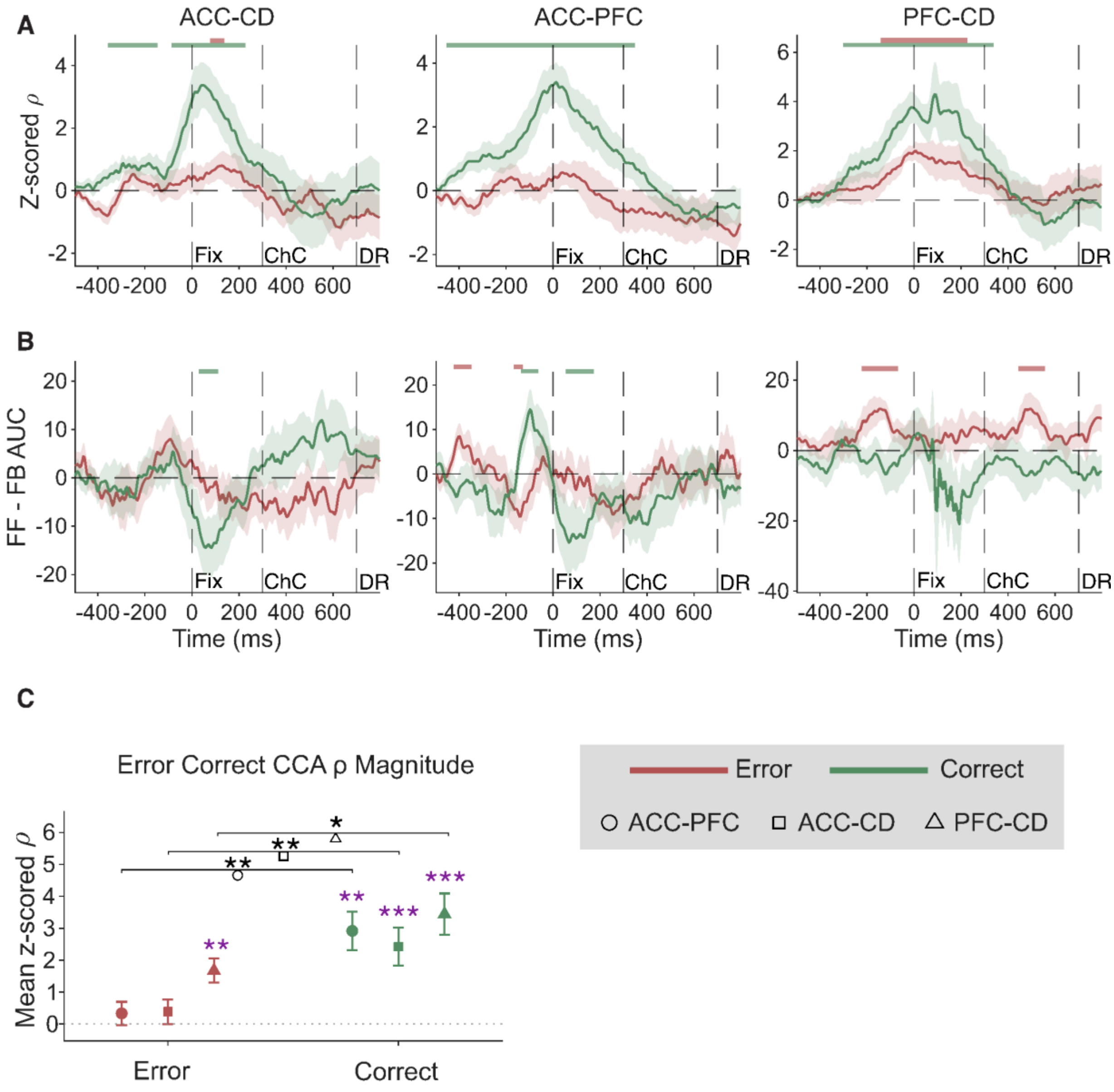


**Fig. S7** (**A**) Z-scored CCA $\rho$ from averaged time-lagged CCA across area pairs for correct trials and error trials with shading from s.e.m. Green and red bars indicate above baseline inter-areal connectivity from cluster permutation tests. (**B**) Directionality for each area pair calculated from subtracting averaged positive time-lagged time course from the negative time-lagged time course with green and red bars indicating greater than baseline deviations in feedforward or feedback dynamics. (**C**) Average peak magnitude plus s.e.m. in the $\pm 100$ ms around fixation. Purple stars indicate Holm-adjusted sign flip test around zero while black stars indicate two sided Holm-adjusted between conditions permutation tests (* $p < .05$, ** $p < .01$, *** $p < .001$ for all tests). For all area pairs, the magnitude of CCA $\rho$ peaks was higher during correct versus error trials and the PFC-CD was the only pair to have a significant peak during error trials.

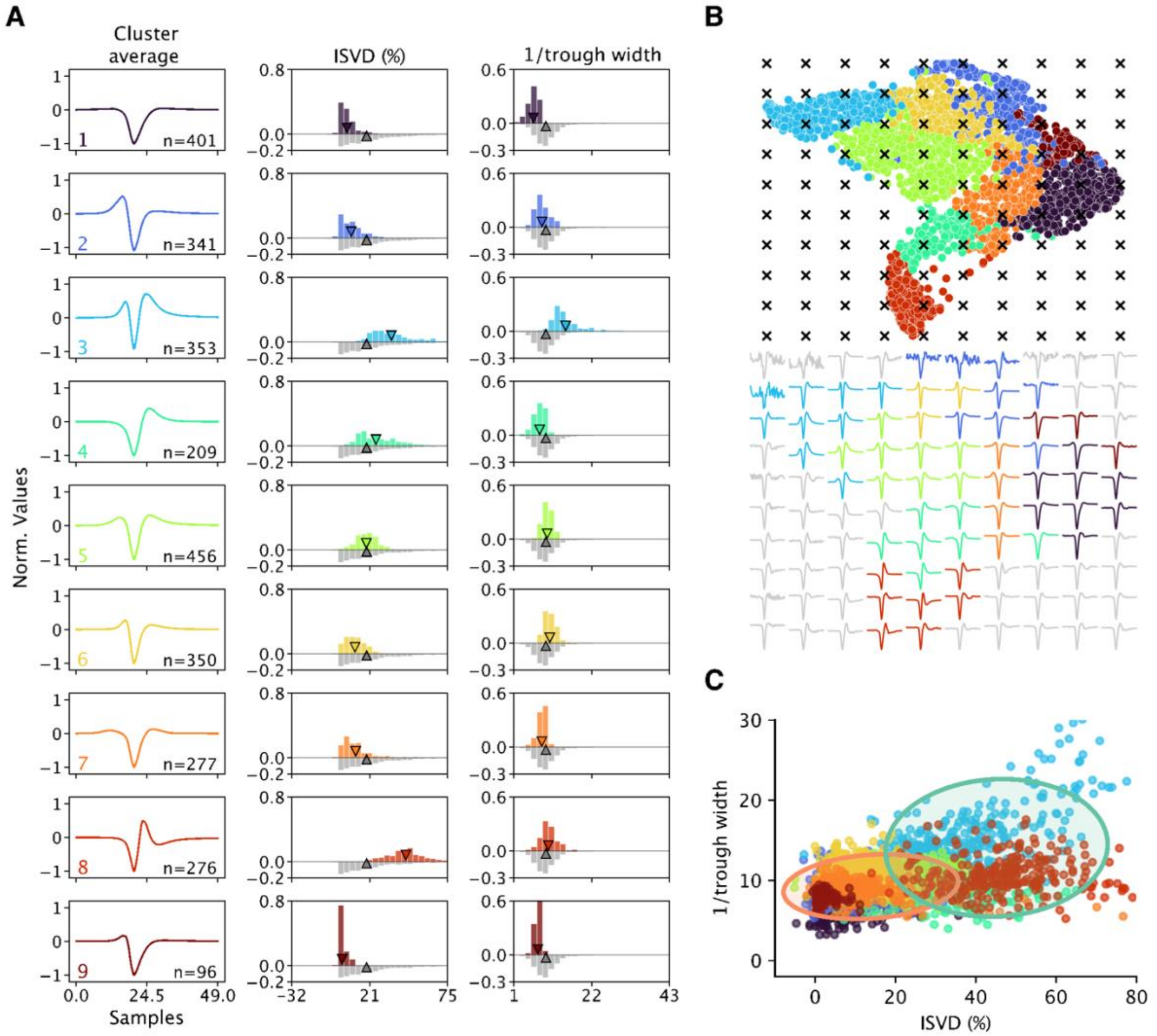


**Fig. S8** (**A**) Visualization of average cluster waveform for WaveMAP sorted neurons in the CD along with the initial slope to valley decay percentage (ISVD %) and inverse trough width distributions for neurons in that cluster. (**B**) U-MAP cluster projected onto the first two U-MAP dimensions along with an inverse transform to visualize the shape of waveforms at each point in the low dimensional U-MAP space. (**C**) Two Gaussian mixture model fit onto a scatter plot of neurons with inverse trough width and ISVD % as the axes. The Gaussians show that seven out of nine clusters are captured by a gaussian with low inverse trough width and ISVD % while two clusters, clusters three and eight are captured by a gaussian with high ISVD % and inverse trough width values, identifying two clusters of putative narrow spiking interneurons.

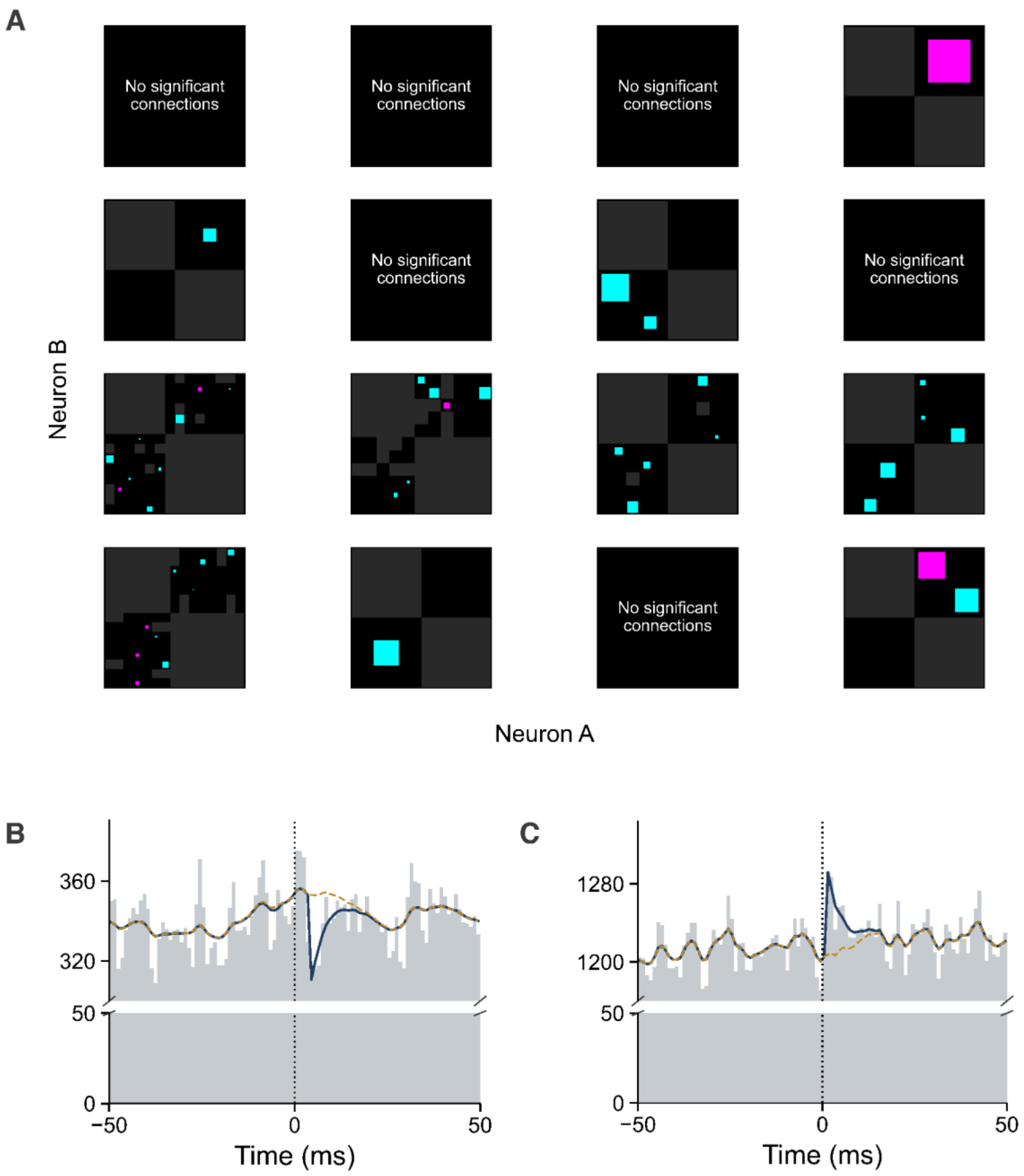


**Fig. S9** (**A**) Connectivity of significant neuronal influences between PFC-CD neurons in both directions from GLMCC analysis across all analyzed sessions with PFC and CD recordings. There were 17 identified (put.) inhibitory influences and 4 identified (put.) excitatory connections (cyan and magenta squares respectively). (**B**) Example inhibitory cross correlogram with directionality from PFC to CD showing a suppression of the CD neuron ~4 ms after the PFC neuron fired. The yellow dotted line represents the fitted baseline while the dark blue solid line represents the GLM fit. (**C**) Same as above but for an example excitatory PFC to CD connection.

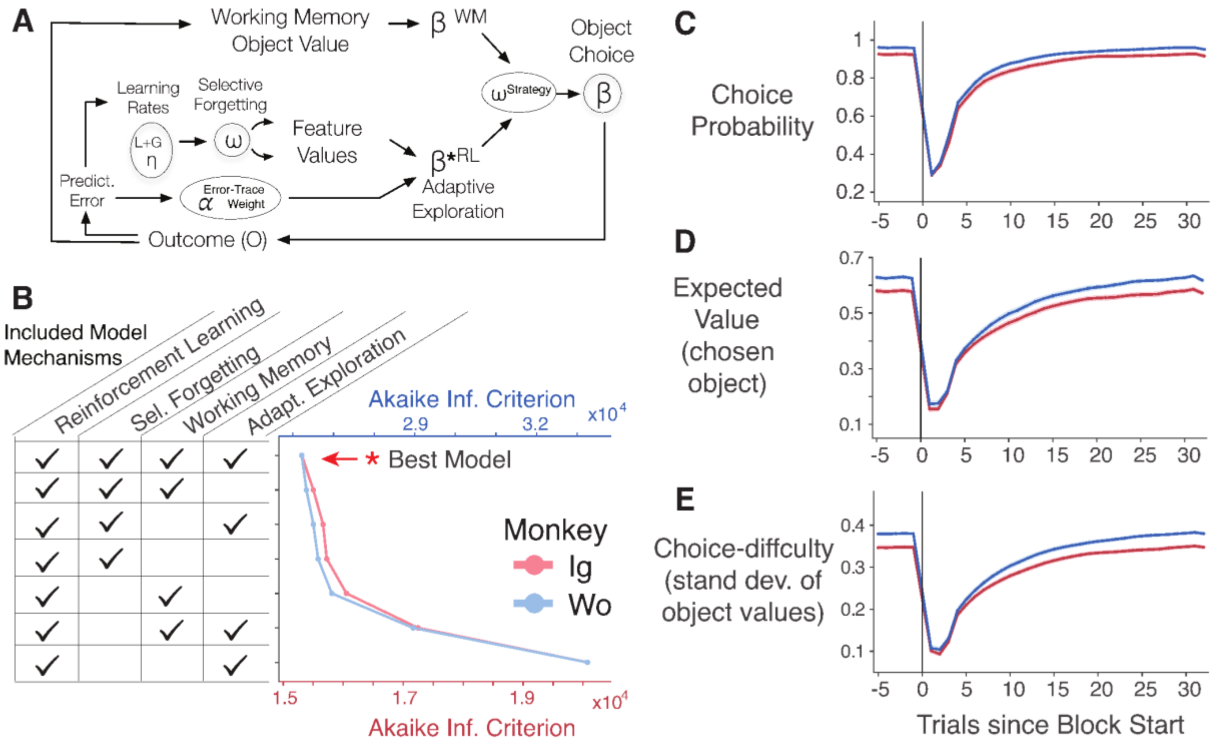


**Fig. S10** (**A**) Schematic of an augmented Rescorla Wagner reinforcement learning model with additional mechanisms for selective forgetting, working memory, and adaptive exploration. (**B**) The best model was chosen based on minimization of Akaike Information Criterion and was found to be the model that included all mechanisms for both monkeys. (**C**) The model-derived probability of choosing the target stimulus plotted across trials from the start of the block for both monkeys. (**D**) Curves of the model derived expected value of the object which was chose. (**E**) The choice difficulty was represented by the standard deviation of the values of all objects with harder choices characterized by smaller differences between object values and easier choices by a larger standard deviation between the rewarded object and the distractors.
